# Musical Training Increases Robustness of Sequence Learning and Increases Anticipatory Responses During Sequence Learning

**DOI:** 10.64898/2025.12.17.695034

**Authors:** Li-Ann Leow, Jarrad Lum, Sara Johnson, Emily Corti, Welber Marinovic

## Abstract

Musicians demonstrate advantages in acquiring motor sequences, showing faster learning and better explicit sequence knowledge than non-musicians. However, it is unclear whether this advantage extends beyond acquisition to the consolidation phase, which is when newly learned skills stabilize and become resistant to interference. Additionally, while interference from executing competing motor tasks is well-established, less is known about whether purely sensory information presented after learning can disrupt consolidation of a bimodal motor sequence.

We investigated how post-acquisition sensory interference affects performance of a learned audio-visual sequence, and whether musical training moderates this vulnerability. Participants first learned an explicit sequence in a serial reaction time task using synchronous, informative audio-visual cues. After a brief consolidation period, they were randomly assigned to one of four observational conditions that manipulated the relationship between auditory and visual streams. Motor performance was then reassessed.

Post-acquisition sensory interference impaired subsequent motor performance, but this effect was modality-specific: it was driven primarily by manipulations to the task-relevant visual stream, while auditory interference alone had no credible effect. Distributional analysis revealed that learning involved a strategic shift from reactive to anticipatory responding. Critically, participants with musical training showed an earlier and consistently higher reliance on anticipatory responses than those without, demonstrating a more rapid adoption of predictive motor control.

These findings demonstrate that newly formed sensorimotor memories are selectively vulnerable to interference in task-relevant modalities. Furthermore, our work provides a candidate mechanistic account for the musician advantage in sequence learning, linking it to faster development of predictive motor strategies during consolidation.

## Introduction

Our memories are typically multisensory, combining sensory experiences encoded by specialised areas of our brains. For example, imagine watching a pianist perform. You see a finger pressing down a key, an action that is immediately followed by the characteristic sound that key makes. Once you learn the association between keys and sounds, you can predict one from the other: what key produced the sound, and what sound that key will produce. Through this learning process, experienced musicians can guide sequences of actions based on the predictions of sensory inputs (Chen et al., 2008a, 2008b, 2009; Chen et al., 2006; Matsushita et al., 2021; Zatorre et al., 2007). Similarly, in experimental studies of sequence learning, auditory stimuli can be used as a cue that predicts the required action (Han et al., 2023; Leow et al., 2025), or as a form of feedback after the motor action (Hoffmann et al., 2001; Stocker & Hoffmann, 2004; Stocker et al., 2003).

Musical training requires acquisition of spatial-temporal sequences that are of far longer lengths than the 8 to 20 element sequences that are used in behavioural paradigms used to quantify sequence learning, such as in the serial reaction time task (SRTT) (Janata & Grafton, 2003). Perhaps unsurprisingly, several studies demonstrate a musician advantage in acquisition of sequences (Anaya et al., 2017; Palmer, 2005; Sobierajewicz et al., 2018; Tierney et al., 2008). Specifically, musical training is associated with a faster rate of training-related reduction in reaction times during sequence learning (Palmer, 2005; Schwizer Ashkenazi et al., 2022) or reduced errors in explicitly recalling previously learnt sequences (Sobierajewicz et al., 2018; Tierney et al., 2008). For example, one study showed that children with musical training showed an advantage in acquiring explicit knowledge of auditory sequences, reproducing more items from recently executed auditory sequences compared to children with similar volume of gymnastics training or video-game training (Tierney et al., 2008). Indeed, some studies suggest a broad advantage in acquiring explicit sequence knowledge, even when sequences were presented only in the visual modality (Anaya et al., 2017; Pau et al., 2013; Schwizer Ashkenazi et al., 2022), when sequences were executed via eye movements (Schwizer Ashkenazi et al., 2022). By contrast, it is not clear if music training improves implicit learning processes, as previous studies show mixed results (Francois & Schön, 2011; Loui et al., 2010; Romano Bergstrom et al., 2012; Thorpe et al., 2020). Musical training might thus augment the capacity to acquire explicit knowledge of sequences.

Whilst there is an increasing body of work demonstrating that musical training can improve the acquisition of explicit knowledge of sequences, less is known about how the effects of music training on learning extends beyond the acquisition and to the consolidation phase. Specifically, during consolidation, the passage of time after initial acquisition increases the robustness of memories, helping newly acquired memories become more resistant to interference and more reliably retrieved (McGaugh, 2000; Wamsley, 2022). A large body of work demonstrates that the passage of time after training can consolidate motor memories in sequence learning tasks (e.g., Walker et al., 2003; Wang et al., 2024) (for a review, see Wamsley, 2022). Although this consolidation has previously been assumed to require the passage of hours after learning, newer evidence demonstrates consolidation at shorter timescales of minutes after initial learning (Wamsley, 2022), whilst the existence of consolidation at timescales of seconds (Bönstrup et al., 2019) is controversial (Das et al., 2025). During the consolidation timeframe, sequence memories are initially labile, as they can be disrupted by learning a competing motor sequence (Robertson, 2012), and may also be either enhanced or disrupted through the simple observation of congruent or incongruent actions (Kelly et al., 2003). However, such sequence memories become increasingly robust with increasing time after learning (Robertson, 2012). It is unclear if musical training alters the consolidation of sequence learning, as there is currently no direct evidence addressing this question. It is possible that musicians show enhanced consolidation of learning, as musical training is associated with more transfer of learning from one auditory-visual sequence to a second auditory-visual sequence (Palmer, 2005). Alternatively, musical training might increase the rigidity of sequence learning, as extensive musical training can reduce flexibility in acquisition of new auditory sequences when there are counterintuitive mappings between the response and the auditory feedback (i.e., where higher-pitched feedback is associated with keys on the left) (Pfordresher & Chow, 2019).

In the present study, we investigated whether musical training alters the consolidation of acquired sequence memories. Specifically, we asked whether musical training renders motor memories more robust against interference after a post-training consolidation phase. To test consolidation of the sequence learning, we first had participants complete an acquisition phase, where they acquired a 10-item motor sequence that was cued by synchronous auditory-visual stimuli, to establish a strong initial learning effect, similar to our earlier work (Leow et al., 2025). After a 2-minute consolidation break, we then interfered with this learning by having participants learn a new, interfering sequence via observation alone. As some previous studies demonstrated that music training has modality-specific effects on sequence learning, where there is a selective benefit for auditory and not visual sequence learning (Tierney et al., 2008), we manipulated the modality of the new, interfering sequence. Participants were assigned to one of four interference conditions: (1) a **Visual Interference** group observed a new visual sequence paired with the original auditory sequence; (2) an **Auditory Interference** group observed the original visual sequence with a new auditory sequence (i.e., a new mapping of sounds to keypresses); (3) an **Audio-visual Interference** group observed a new visual sequence paired with a new auditory sequence; and (4) a **No Interference** control group observed the exact same audio-visual sequence from the acquisition phase. This interference phase required no motor responses from participants, but they were told that they would have to report that sequence later.

While sequence learning is often assessed through reductions in averaged reaction times (RTs), this can obscure underlying cognitive changes. For example, performance improvements may reflect not just faster motor execution, but also a shift from reactive to predictive control during sequence learning. Supporting this, oculomotor tracking studies have shown that anticipatory eye movements offer a rich, real-time index of multiple, simultaneous learning processes that are not captured by averaged reaction times alone (Tal et al., 2021). Similarly, studies that tightly control movement preparation time in sequence learning tasks suggest that movements during sequence learning can be pre-planned at least 3 movements in advance (Ariani & Diedrichsen, 2019). Increasing evidence also shows that such movement pre-planning occurs during execution of an ongoing movement (Ariani & Diedrichsen, 2019; Kashefi et al., 2024). To capture these subtler dynamics, we examined the full distribution of reaction times rather than relying solely on mean values. This approach, inspired by recent work demonstrating the complexity of sequential effects, enables identification of distinct cognitive mechanisms that influence different portions of the RT distribution (Voormann & Miller, 2024). Moving beyond average RT is thus essential, as even indirect behavioural markers (such as prediction-driven shifts in response time distribution) might reveal learning dynamics that would otherwise remain hidden.

## Methods

### Participants

A total of 151 undergraduate psychology students were recruited from the Curtin University Psychology SONA Participant Pool and participated in exchange for course credit. The description of the study on the SONA participant pool made no mention of musical training. Participants were asked about musical training only at the end of the study session. The sample included 115 females, 34 males, and 2 participants who identified as other, with a mean age of 20.5 years (SD = 2.74). Regarding experience with musical training, 74 participants (49.0%) reported having no experience with formal musical training, while the remaining 77 participants (51.0%) had at least some experience with formal musical training. All participants had normal or corrected-to-normal vision and hearing, and no known cognitive or neurological conditions that could have affected their performance. The study was approved by the Curtin University Human Research Ethics Committee (HRE2018-0257), and all participants provided informed consent prior to taking part in the study.

### Sample size

We did not conduct a formal a priori power analysis for the present study. Instead, we aimed to recruit a sample size comparable to our previous work on bimodal sequence learning. The present study includes approximately 38 participants per condition, which exceeds the sample size of ∼32 participants per group used in Leow et al. (2025). We reasoned that this sample size would provide adequate power to detect the effects of interest related to sensory interference.

### Apparatus and Materials

Visual and auditory stimuli were presented using Inquisit 6 software (Version 6.6.0). Participants were seated in front of a 24-inch computer monitor (1920×1200, 60Hz) and used a standard QWERTY keyboard for responses. Sounds were presented through Corsair HS50 stereo headphones at a fixed intensity of 65dBa.

### Task and Stimuli

Participants performed a multisensory serial reaction time (SRT) task. They were seated in front of a computer and placed the middle and index fingers of their left hand on the ‘V’ and ‘B’ keys, and the index and middle fingers of their right hand on the ‘N’ and ‘M’ keys, respectively. The screen displayed four grey squares in a horizontal row, which mapped spatially from left to right to the four response keys. On each trial, one square changed colour for 100 ms. This colour change occurred synchronously with a 100 ms auditory tone. The auditory stimuli were four pure tones (C4, D4, E4, and F4) that were mapped informatively to the four visual locations, creating a consistent audio-visual pairing during the acquisition phase (detailed later). Participants were instructed to respond as quickly and accurately as possible by pressing the key corresponding to the square that changed colour. Following a correct response, the trial ended, and a 500 ms inter-trial interval occurred before the next trial began (i.e., there was a 500 ms response-stimulus interval). However, in the interference phase, the stimulus-response mapping was altered as per the task condition.

### Procedure

The task comprised three main phases: acquisition, interference, and revision (see Figure 1B for an overview). First, participants were familiarized with the task by completing 50 practice trials with random stimuli. They then began the Acquisition Phase (Blocks 1-5), which consisted of five blocks of 100 trials each (each block = 10 repetitions of a 10-item sequence per block): this resulted in a total of 500 trials for the acquisition phase. In this phase, all participants learned the same sequence with synchronous and informative audio-visual stimuli (Figure 1A). Following acquisition, a two-minute Consolidation Break was administered, during which participants did not perform any task: this timeframe is sufficient to allow rapid offline consolidation for sequence learning (Bönstrup et al., 2019). Next, in the Interference Phase, participants were randomly assigned to one of four interference conditions. This phase consisted of a single block (100 trials, or 10 repetitions of a new or the original 10-item sequence) that was purely observational, requiring no motor responses, similar to previous work by Howard and colleagues (Howard et al., 1992). The four conditions were: (1) a Visual Interference condition, in which participants passively observed a new visual sequence paired with the original auditory sounds (the original auditory cue order and auditory-response mapping was maintained, but the responses were now mapped to spatially different visual cues); (2) an Auditory Interference condition, in which participants passively observed the original visual sequence with new auditory sounds (the original visual sequence order was maintained and the visual cue-response mapping was maintained, but the auditory-response mapping was now altered); (3) an Audio-visual Interference condition, in which participants observed a new visual sequence paired with new sounds (both the visual and auditory cue order was altered); and (4) a No Interference control condition: this group observed the exact same audio-visual sequence from the acquisition phase. Participants were instructed that they would have to report the sequence they observed in this phase later.

**Figure 1.**
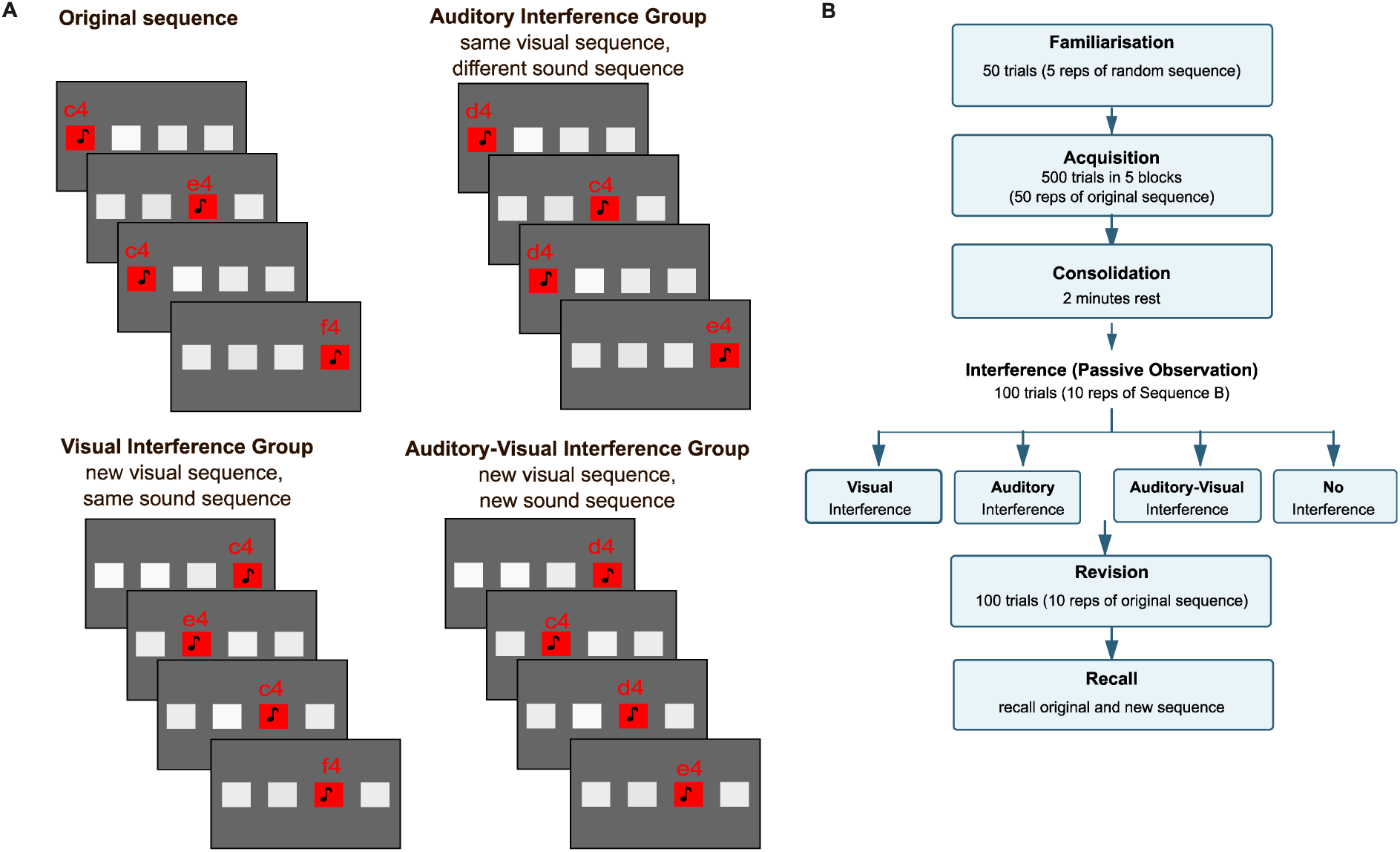
(A) **Experimental design showing the four interference conditions**. Participants responded to visual cues (red squares) paired with corresponding auditory tones using keyboard presses with both hands. During acquisition, the original sequence (1,3,1,4,3,2,4,2,3,1) had each visual cue position informatively paired with a specific auditory tone (1=C4, 2=D4, 3=E4, 4=F4). Following acquisition, participants were assigned to one of four passive observation interference conditions that manipulated the congruence between visual and auditory cues. In the **Visual Interference group** (bottom left), participants passively observed a new visual sequence while the original auditory sequence was maintained. In the **Auditory Interference group** (top right), participants observed the original visual sequence paired with new auditory tones; the visual sequence order and visual cue-response mappings were preserved, but auditory-response mappings were altered. In the **Auditory-Visual Interference group** (bottom right), both visual and auditory sequences were changed, with responses mapped to new visual positions and paired with new auditory tones. A **No Interference control group** passively observed the original audiovisual sequence from the acquisition phase. (B) Schematic of the experimental procedure. Following familiarization (50 trials), participants completed an Acquisition phase (500 trials across 5 blocks) learning a 10-item sequence with synchronous audio-visual stimuli. After a 2-minute Consolidation break, participants were randomly assigned to one of four Interference conditions (100 trials of passive observation): Visual Interference (new visual sequence with original sounds), Auditory Interference (original visual sequence with new sounds), Auditory-Visual Interference (new visual and auditory sequences), or No Interference (original sequence). Subsequently, participants performed a Revision phase (100 trials) with the original sequence, followed by a Recall phase testing explicit memory for both sequences.

**Figure 2.**
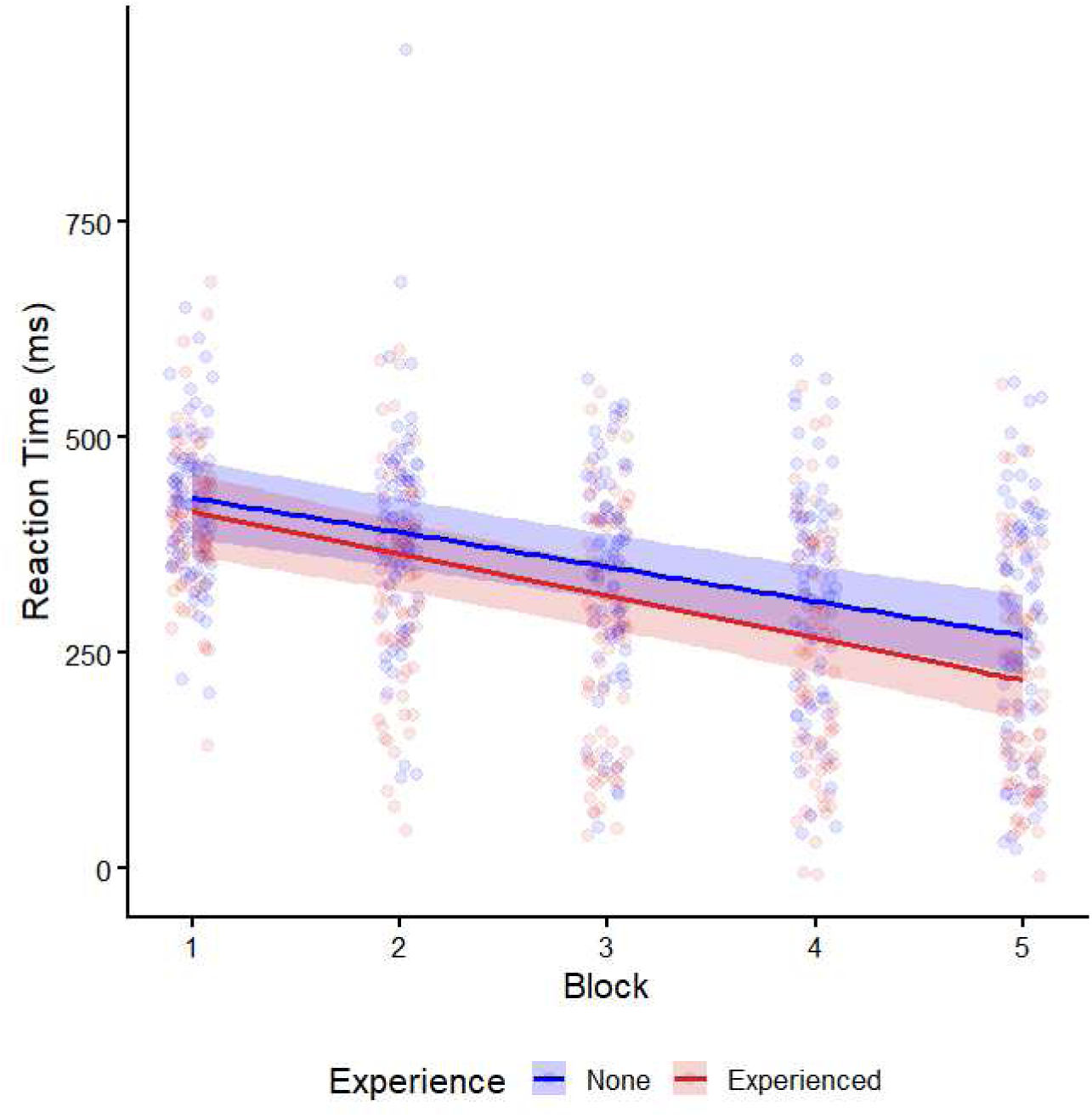
Analysis of the Learning Phase and Performance at the End of Acquisition. Conditional effects from the Bayesian model showing the estimated learning rate for participants with None or some Musical Experience across the five acquisition blocks. Solid lines represent the posterior mean estimates for Reaction Time (ms), and the shaded ribbons represent the 95% credible intervals.

Subsequently, in the Revision Phase (Block 6), all participants performed one block of 100 trials of the original acquisition sequence to measure the effect of the observational interference on their motor performance. Finally, in the Recall phase, participants were asked to explicitly recall the original sequence as well as the interference sequence, using mouse clicks.

### Data Cleaning and Trial Exclusion

Prior to the main analyses, the data was processed to exclude trials with incorrect responses or slow reaction times (RT). From a total of 90,600 trials, responses were removed if they were incorrect or if the RT exceeded a 1000 ms cutoff. This procedure led to the exclusion of 5,530 trials (6.10%) for incorrectness and 1,971 trials (2.18%) for exceeding the RT limit. After accounting for trials that met both criteria, a total of 6,862 unique trials were removed, representing 7.57% of the total data. Consequently, 83,738 trials (92.43%) were retained for the final analyses.

### Data Analysis

All statistical analyses were performed using R (version 4.3.2) with the brms package (Bürkner, 2017) for Bayesian multilevel modelling. To account for the heavy-tailed distribution of reaction times and provide robustness against outliers, we utilised a Student-t likelihood. Instead of aggregating trials into block means, our primary analyses utilised trial-level data to preserve the natural variance of motor performance (Vehtari et al., 2021).

Following best practice in Bayesian modelling (Kruschke, 2021), we specified weakly informative, data-aware priors to improve estimation stability and guard against overfitting.

For continuous outcomes, intercepts received normal priors centred on the observed mean; slopes and interaction terms received normal priors centred on zero. We employed a maximal random-effects structure including random intercepts and random slopes for the temporal predictors (Block or Phase) per subject (Barr et al., 2013). To ensure numerical stability and improve MCMC sampling efficiency, we used a non-centred parameterization (decoupling the correlation between intercepts and slopes) (Vehtari et al., 2021).

Following the recommendations of Vehtari et al. (2021), we verified that all rank-normalized split-Rhat values were < 1.03. We estimated the Monte Carlo Standard Error (MCSE) to ensure that the precision of our posterior estimates was sufficient for the nature of our data (reaction time). We prioritised Tail-ESS values above 400 to ensure the stability of the reported 95% Credible Intervals (CrI). For all models, we report the median of the posterior distribution and the 95% CrI.

The acquisition phase (Blocks 1–5) was analysed with a model including Block (mean-centred), Condition, and Musical Experience as interacting predictors. For the interference analysis, we utilised a three-way interaction model (RT ∼ Phase * Condition * Musical Experience) comparing the final acquisition block (Block 5) to the post-interference block (Block 6). This interaction approach effectively quantifies the interference effect on learning while accounting for individual baseline performance at the end of acquisition.

A second set of analyses targeted explicit memory measures collected at the end of the experiment. We fitted Bayesian models (Student-t likelihood with robust priors) to explicit memory scores for the practiced and interference sequences. Finally, we examined distributional aspects of RTs. A two-component normal mixture model on log-transformed latencies provided a data-driven threshold separating fast, anticipatory responses from slower, reactive responses. Using this threshold, we computed (1) the proportion of anticipatory responses per block, analysed with a Bayesian linear mixed model, and (2) a binary bimodal classification for each participant and block, analysed with a Bernoulli Bayesian GLMM. This allowed us to characterise how the prevalence of anticipatory and bimodal response patterns evolved across practice and differed by musical experience.

## Results

### Acquisition Phase: Blocks 1–5

During the acquisition phase, participants in all groups experienced identical synchronous and informative audio-visual stimuli. Still, we sought to test for pre-existing group differences during this acquisition phase prior to the primary intervention in the Interference phase. Thus, we employed a maximal Bayesian linear mixed-effects model. While some marginal parameters exhibited R-hat values up to 1.03, the Monte Carlo Standard Error (MCSE) for primary effects remained below 0.5 ms (representing <2% of the total effect magnitude), and Tail-ESS values for all critical interaction terms exceeded the recommended threshold of 400, confirming the precision and stability of the reported 95% CrI. Our results revealed a robust main effect of Practice, with reaction times (RT) decreasing by an average of 39.83 ms per block (95% CrI [−54.10, −25.87]; MCSE = 0.48 ms). Critically, there were no credible interactions between Block and the eventually assigned interference conditions (all 95% CrIs for Block x Condition terms crossed zero), confirming that all experimental groups were balanced in their learning capacity prior to the intervention. In addition, while the mean estimate for the Block × Musical Experience interaction suggested that experienced musicians learned at a faster rate than non-musicians (Estimate = −8.77 ms), the 95% CrI for this effect included zero ([−26.86, 10.26]), precluding a conclusion of a credible difference at this threshold. This suggests that while musical experience may offer a slight advantage, individual trial-level variability was substantial enough to obscure a robust group-level learning difference.

### Analysis of Response Time Distributions During Acquisition

An inspection of the trial-level distribution of the whole dataset suggested a potentially complex underlying structure. Specifically, the distribution of reaction times appeared to be bimodal, with responses clustering into a fast, anticipatory mode and a slower, reactive mode (Figure 3A). To gain a more mechanistic understanding of how learning unfolded, we conducted a subsequent analysis to explicitly model these two response components.

**Figure 3.**
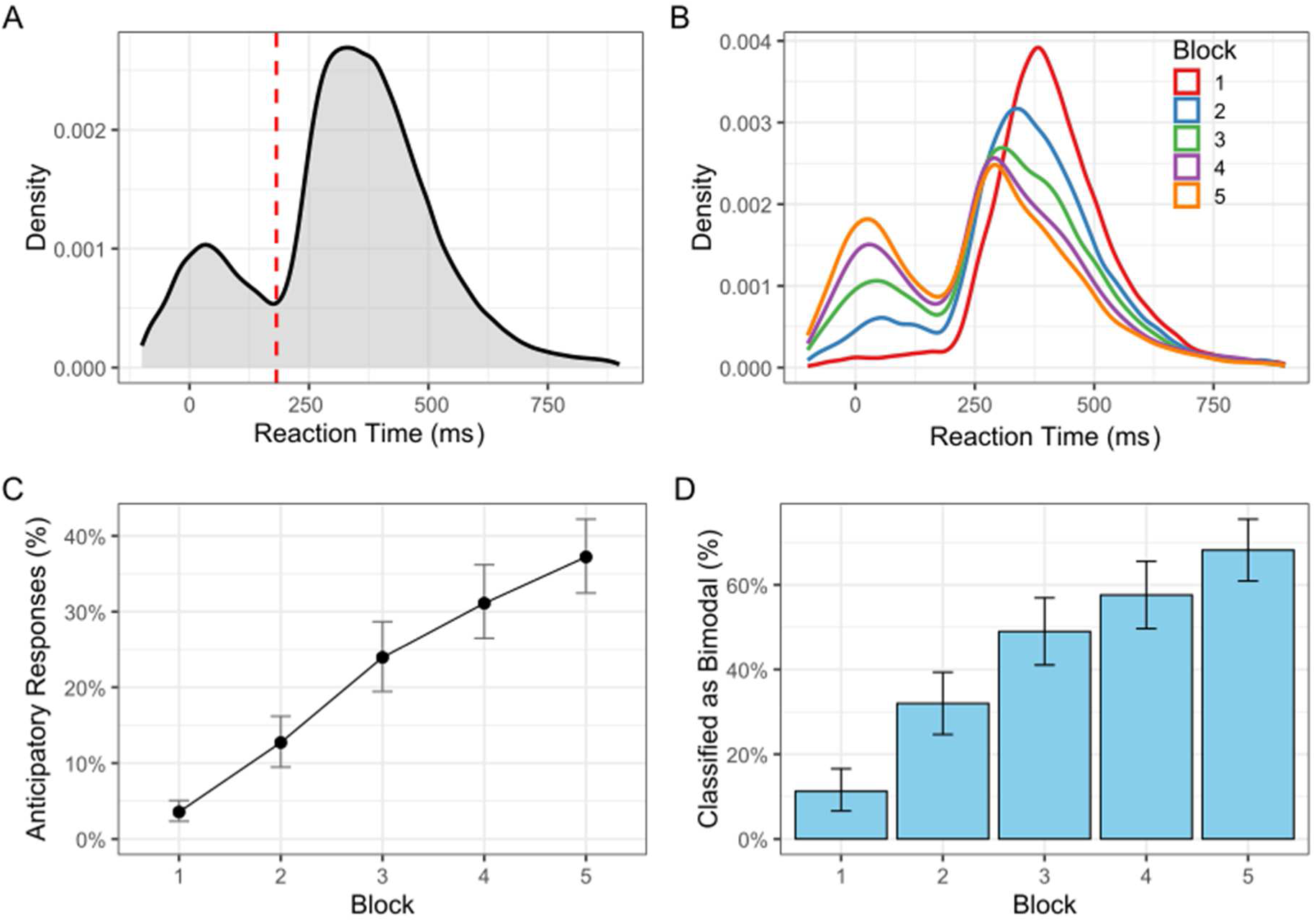
Emergence of a Bimodal Distribution in Reaction Times During Learning. (**A**) The density plot of all reaction times (RTs) from the acquisition phase, collapsed across all blocks and participants. The plot clearly shows a bimodal distribution, suggesting the presence of two distinct response types: a fast, anticipatory mode and a slower, reactive mode. (**B**) The density plot of RTs separated by training Block. This panel illustrates the dynamic shift in response strategy during learning. In Block 1 (red line), the distribution is largely unimodal and centred on slower, reactive responses. As practice progresses to Block 5 (orange line), a distinct peak of fast, anticipatory responses emerges and grows, indicating an increasing reliance on predictive motor control. (**C**) The plot shows the mean proportion of anticipatory responses across the five acquisition blocks. Points represent the mean, and the vertical bars represent the 95% bootstrapped confidence intervals. (**D**) The bars represent the mean proportion of subjects classified as “bimodal” in each of the five acquisition blocks. Error bars represent the 95% bootstrapped confidence intervals.

### Practice Increases Anticipatory Responding During Acquisition of Sequences

To quantify the shift from reactive to predictive control, we first defined a data-driven threshold to separate the two response modes. A two-component mixture model was fitted to the distribution of all log-transformed reaction times from the acquisition phase, yielding an anticipation threshold of 182.01 ms. This threshold reflects a global mixture solution estimated across participants and acquisition blocks, assuming stable response mode structure across groups and practice. As shown in Figure 3B, the distribution of reaction times clearly shifted across the acquisition phase, with a growing peak of fast, anticipatory responses emerging over time.

To formally test this, we first examined the overall trend across all participants. A Bayesian mixed-effects model revealed a credible posterior effect of practice, showing that the proportion of anticipatory responses (latencies < 182.01 ms) increased significantly across blocks (Median = 0.95, 95% CrI [0.85, 1.05]; max R-hat = 1.001; Bulk ESS ≈ 1121; Tail ESS ≈ 2132; see Figure 3C). Similarly, we classified each participant on a block-by-block basis as either showing a bimodal latency distribution or not. A Bayesian logistic mixed-effects model confirmed that as participants practiced, they were significantly more likely to develop a bimodal response pattern (Median = 2.03, 95% CrI [1.55, 2.62]; max R-hat = 1.001; Bulk ESS ≈ 4079; Tail ESS ≈ 6758; see Figure 3D).

### Music Training Increases Anticipatory Responding During Acquisition of Sequences

To understand the source of the different learning rates observed in our initial analysis, we next examined how music training moderated this strategic shift towards anticipatory responding during acquisition. Following a maximal modelling approach, we included the interaction between Block and Musical Experience to test for differences in both overall levels and rates of strategy adoption.

For the proportion of anticipatory responses, we found a credible posterior main effect of Musical Experience. Participants without music training were significantly less likely to anticipate compared to experienced musicians (Estimate = −1.38, 95% CrI [−2.23, −0.51]). However, the interaction between Block and Experience was not credible (Estimate = 0.03, 95% CrI [−0.16, 0.23]), indicating that while musicians maintained a higher baseline of anticipation, the *rate* at which both groups adopted this strategy was statistically similar.

The analysis of bimodal reaction time distributions yielded a similar pattern. We observed reliable posterior evidence for a main effect of Musical Experience, with non-musicians being significantly less likely to be classified as bimodal responders throughout acquisition (Estimate = −1.53, 95% CrI [−2.79, −0.28]). As with the proportion model, the interaction term for bimodality was not credible (Estimate = 0.00, 95% CrI [−0.64, 0.66]). These results suggest that musical training provides a robust, overall advantage in the ability to generate anticipatory motor actions, though the trajectory of this strategic shift with practice is consistent across experience levels (Figure 4B).

**Figure 4.**
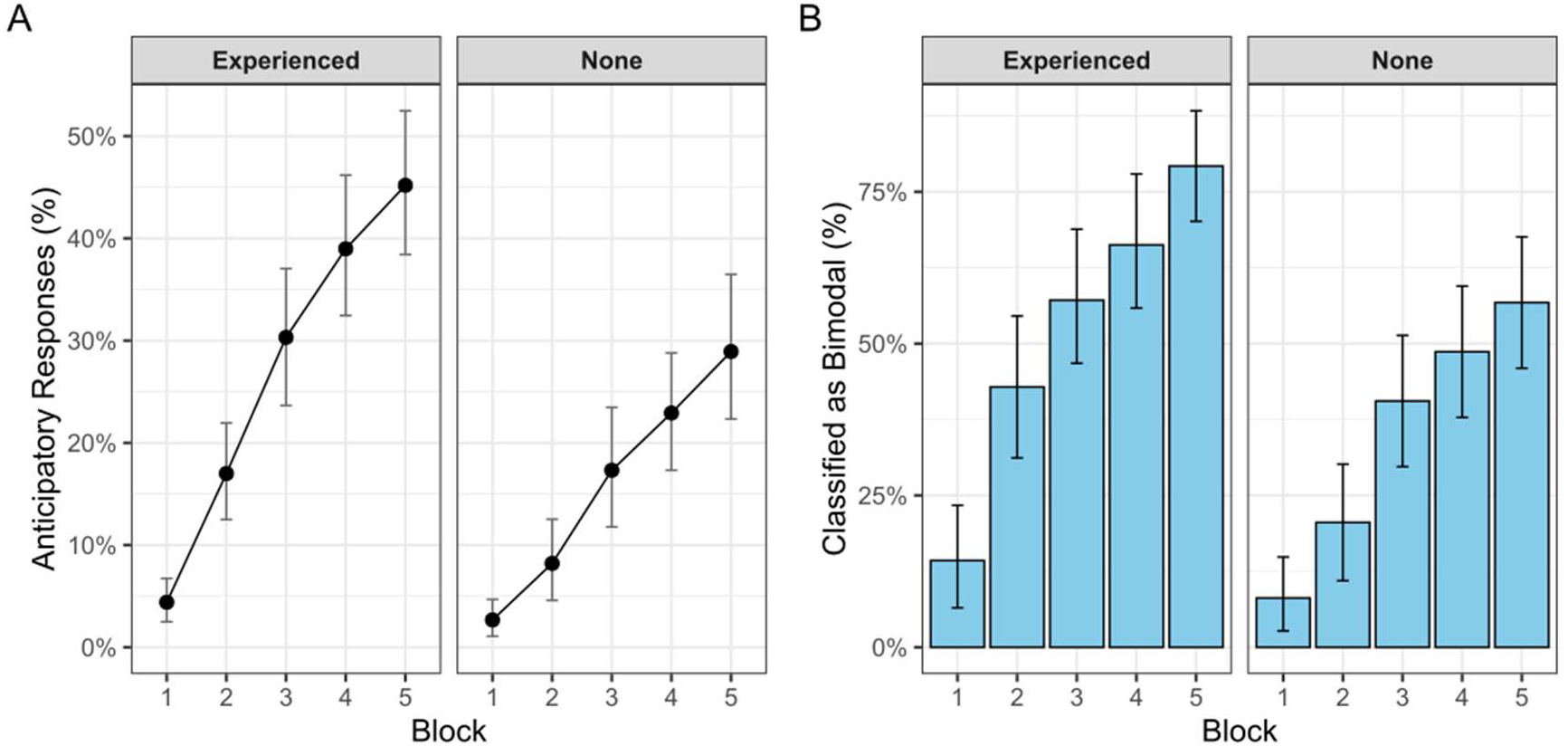
Moderating Effect of Experience with Musical Training on the Emergence of Anticipatory Responses During Acquisition. **(A)** The proportion of anticipatory responses across the five acquisition blocks, plotted separately for participants with (“Experienced”) and without (“None”) musical training. Points are the means and error bars represent 95% bootstrapped confidence intervals. **(B)** The proportion of participants classified as having a bimodal RT distribution in each block, separated by experience with musical training. Bars represent the mean proportion and error bars represent the 95% bootstrapped confidence intervals.

### Interference from Observing a New Visual Sequence

We next tested how observing a new sequence interfered with the newly formed motor memory by comparing performance in the final acquisition block (Block 5) to the post-interference block (Block 6). To identify modality-specific disruption, we utilised a Bayesian multilevel interaction model (RT ∼ Phase x Condition x Musical Experience) with a maximal random-effects structure. Although some marginal parameters exhibited R-hat values up to 1.03, the MCSE for the critical interaction terms remained low (∼1.2–1.8 ms), and Tail-ESS values for these parameters exceeded the recommended threshold of 400.

The analysis revealed a significant Phase x Condition interaction, indicating that the effect of the interference on previous learning was modality-dependent. In the reference group (Auditory Interference), participants continued to demonstrate robust performance gains, with reaction times (RT) decreasing by an average of 82.24 ms (95% CrI [−116.01, −47.09]; MCSE = 1.01 ms). However, this learning trajectory was credibly hindered in the Bimodal audio-visual (Estimate = +70.31 ms, 95% CrI [21.62, 120.04]; MCSE = 1.23 ms) and Visual (Estimate = +60.19 ms, 95% CrI [12.76, 109.06]; MCSE = 1.36 ms) interference groups. In these conditions, the positive interaction terms effectively counteracted the expected practice gains (resulting in net RT changes of only −11.93 ms and −22.05 ms, respectively as shown in Figure 5). In contrast, the No Interference group did not credibly differ from the auditory reference group (Estimate = +15.78 ms, 95% CrI [−30.17, 61.25]).

**Figure 5.**
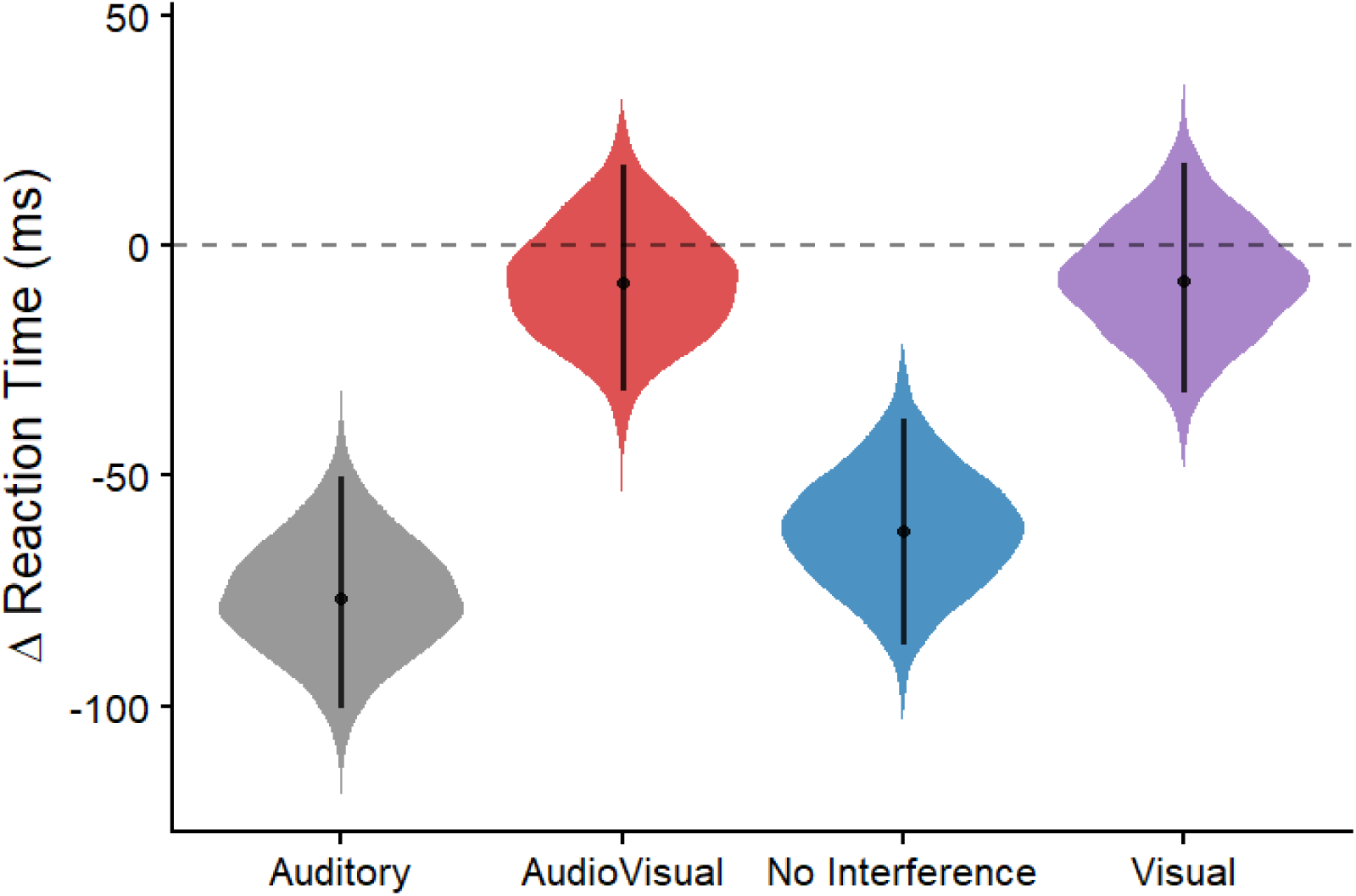
Posterior Distributions of Learning Change (Delta RT). Vertical violin plots show the density of the posterior distribution for the change in reaction time between Block 5 and Block 6 for each condition. Negative values indicate reductions in reaction time. While participants who observed an Auditory or No Interference sequence showed continued speed gains, those in the Visual and Bimodal groups showed a credible slowing of learning. Points indicate the posterior median; bars indicate the 95% HPDI. Estimates are derived from a trial-level interaction model accounting for individual learning trajectories.

Regarding the role of expertise, we found no credible evidence that musical training moderated the impact of interference. While musicians were faster overall (Estimate = −51.32 ms), all two-way interactions involving musical experience (e.g., Phase x Musical Experienced) and all three-way interactions (e.g., Phase x Condition x Musical Experience) yielded 95% CrIs that crossed zero. These results indicate that while musical experience may provide an advantage in raw speed and acquisition rate, it does not offer a significant protection against the disruption caused by task-relevant visual-spatial interference.

Consistent with our acquisition phase findings, these results suggest that interference was primarily driven by the manipulation of the task-relevant visual stream, regardless of prior musical training.

## Explicit Memory

### Better Explicit Memory for the Practiced (Acquisition) Sequence With Music Training

To assess participants’ explicit memory for the sequence practiced in the Acquisition Phase, we quantified recall in the Revision Phase using mouse-click sequences. We fitted a Bayesian linear model (Student-t likelihood) with condition and musical experience as interacting predictors. The analysis did not reveal a credible interaction between the interference condition and musical experience; participants with and without musical training demonstrated broadly similar recall scores (Figure 6, left panels).

**Figure 6.**
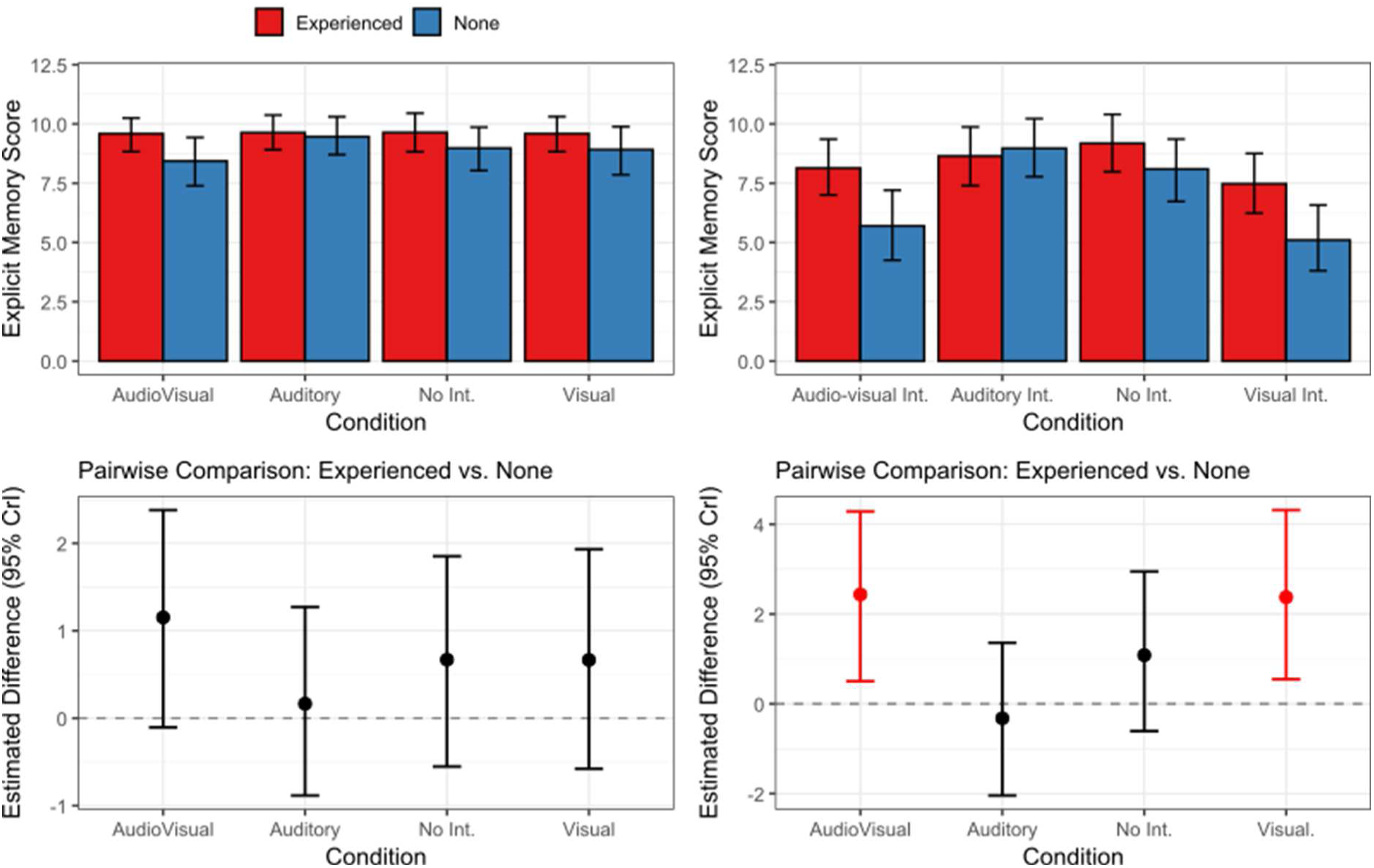
Bayesian Analysis of Explicit Memory Scores. (Left) Memory for the Practiced (Acquisition) Sequence. The top panel displays the estimated marginal mean recall scores for each condition, separated by musical experience. Bars represent the posterior median estimates, and the vertical error bars represent the 95% CrIs. The bottom panel shows the pairwise comparisons of the estimated difference between experienced and non-experienced participants within each condition. Points represent the median of the posterior distribution for the estimated difference, and the vertical bars show the 95% CrIs. **(Right) Memory for the Observed (Interference) Sequence.** The top panel displays the estimated marginal mean recall scores for the second, observed sequence. The bottom panel shows the corresponding pairwise comparisons between experience groups for each condition. In both comparison plots, intervals not crossing the zero line indicate a credible performance difference between those with experience of music training and those without.

Pairwise comparisons further confirmed the lack of a robust musical training advantage. Median differences between groups were small, and their 95% CrIs all included zero (e.g., Audio-visual Interference interaction: Median = −1.00, 95% CrI [−2.64, 0.63]). Following Vehtari et al. (2021), all posterior diagnostics indicated excellent convergence and stability, with all rank-normalized split-R-hat < 1.00 and Tail-ESS values for all fixed effects exceeding 6,400. Overall, these results suggest that those with and without formal musical experience had comparable explicit recall of the originally practiced sequence.

### Better Explicit Memory for the Observed (Interference) Sequence With Music Training

We next analysed participants’ memory for the second sequence observed during the manipulation phase. The results pointed towards an interaction between condition and musical experience (Figure 6, right panels). Pairwise comparisons showed that participants experienced with formal music training outperformed those without music training in remembering the new sequence when it involved visual changes, with credible evidence for this difference in the Visual Interference group (Median Difference = −2.71, 95% CrI [−5.22, −0.24]) and the Audio-visual Interference group (Median Difference = −2.77, 95% CrI [−5.27, −0.28]). As expected, there was no musical experience advantage in the No Interference condition (Median Difference = −1.43, 95% CrI [−3.88, 0.96]). All posterior diagnostics confirmed excellent mixing and convergence (all R-hat ≤ 1.01, all Tail-ESS > 5,400). This indicates that non-musicians struggled more to encode the new, interfering visual sequence itself.

## Discussion

The present study explored the effect of musical training on the capacity to acquire and consolidate distinct sequence memories, and characterised how musical training alters anticipatory response patterns in sequence learning. We found that musical training not only increased the capacity to acquire explicit knowledge of the practiced sequence during the acquisition blocks, but also enhanced strategic and explicit aspects of sequence learning. However, this advantage did not confer protection against interference at the level of motor execution speed of the practiced sequence from interference resulting from observing and learning a distinct visual sequence in the interference block. Furthermore, the response times distributions during the acquisition block also differed depending on musical training. Response times distributions became increasingly bimodal with increasing practice of the sequence in the acquisition block, demonstrating a fast, anticipatory mode and a slower, reactive mode of response. Those with musical training showed an earlier and consistently higher reliance on anticipatory responses than those without, and were also more likely to be classified as bimodal responders than those without musical training. Overall, our findings demonstrate that musical training confers multiple advantages in sequence learning: increasing the robustness of explicit and strategic aspects of sequence learning against interference, increasing explicit recall of the interfering sequence, and increasing predictive control earlier during acquisition of a motor sequence.

### Musical Training Increased the Acquisition and the Robustness of Explicit Sequence Learning

Musical training requires the concurrent acquisition of spatial sequences as well as temporal sequences, where sequences are typically of far longer lengths than the 8 to 20 element sequences that are used in sequence learning tasks like the SRTT (Janata & Grafton, 2003). It is perhaps unsurprising then that musicians show an advantage in sequence learning tasks with auditory-visual sequences (Anaya et al., 2017; Palmer, 2005; Pau et al., 2013), as this constitutes a form of near transfer. Importantly, whilst previous work has demonstrated that musical training improves acquisition of sequences, relatively little has been known about whether musical training alters the *consolidation* of sequence learning, such that memories become more robust against interference. Furthermore, exactly what components of learning are susceptible to interference remain incompletely understood. Previous work has shown that perceptual experience of a sequence is sufficient for acquisition of sequence learning (Howard et al., 1992), although it is less clear if perceptual sequences can also cause interference. We show here that perceptual sequences can cause interference, but not all forms of perceptual stimuli cause interference. Only sequence stimuli presented in the visual modality interfered with previous learning, slowing reaction times when re-executing the original sequence, both in those with and without musical training. Listening to a different auditory sequence did not interfere with previous learning, similar to previous work in expert musicians (Brown & Palmer, 2013). Our results showed that musical training failed to render sequence learning immune from interference from observing a new visual or auditory-visual sequence, slowing reaction times when re-executing the original sequence. What might explain the modality-specificity of this interfering effect? We propose that despite presenting the original sequence with both auditory and visual stimulus information at acquisition, learning occurred predominantly by mapping spatial information of the visual cues to the spatial information of the upcoming action, whereas auditory stimuli served mainly as an accessory stimulus. In other words, learning occurred in a task-relevant domain: where the spatial locations of the cues helped encode the upcoming action. However, the interference effects shown from reaction time decrements in both those with and without musical training contrast with the explicit recall data, where those with musical training were more resilient against interfering effects from the visual sequence when explicitly recalling the initially acquired sequence, as well as the second, interfering sequence.

We note that our task allows concurrent engagement of implicit and explicit learning processes. It remains to be seen if the musical training advantage in both acquiring and consolidating explicit sequence memories here would also be evident for implicit sequence learning processes, as previous findings have been mixed (Francois & Schön, 2011; Loui et al., 2010; Romano Bergstrom et al., 2012; Thorpe et al., 2020). The separable mechanisms of implicit and explicit sequence learning has long been proposed (Reber & Squire, 1998), and evidence suggests that consolidation processes for implicit and explicit sequence learning depend on distinct neural processes (Hirano et al., 2017). For example, perturbational TMS experiments show that offline improvements following the explicit sequence learning depend on the primary motor cortex and not on the inferior parietal lobe, whilst offline improvements following the implicit sequence learning dependent on the inferior parietal lobe but not the primary motor cortex (Breton & Robertson, 2017). It is plausible that the explicit representation formed in our task is less reliant on this low-level motor trace and is more dependent on higher-order cognitive structures, the very networks implicated in long-term musical training (Ihalainen et al., 2023). It remains to be seen if sequence learning paradigms that dissociate the implicit component of learning (e.g., probabilistic serial reaction time tasks, contextual cueing, or statistical learning paradigms) would reveal similar advantages for musicians. Examining these processes more systematically could help clarify whether musical expertise enhances domain-general implicit sensitivity to sequential regularities, or whether the advantage is more specific to explicit strategic learning and higher-order representational processes.

### Prior Musical Training Increased Anticipatory Responding During Sequence Learning

Our analysis of fast, anticipatory reaction times during sequence learning provides a more direct window into the development of predictive processing during sequence learning than standard approaches using mean reaction times. Our analysis supports a longstanding view that a characteristic feature of sequence learning is a shift from a reactive mode of responding to an anticipatory one (Ghilardi et al., 2003). Importantly, participants with musical training showed an earlier shift toward anticipatory responding, supporting the premise that prior expertise can facilitate the development of predictive models for action. Although some previous studies have posited that a shift to anticipatory responding occurs more rapidly in individuals with musical training (Engel et al., 1997; Palmer, 2005)—supporting the idea that prior expertise may facilitate the development of predictive models for action— there is relatively limited direct evidence to substantiate this claim. For example, Engel et al. (1997) demonstrated anticipatory adjustments in skilled pianists’ hand and finger kinematics, but their small sample included a mix of experience levels and did not systematically test training effects; anticipatory modifications were notably task-dependent and variable across both pieces and individuals. Another study found more anticipatory responding and explicit sequence recognition in participants with better aptitude in discriminating musical rhythms, but found mixed evidence for a musician benefit to sequence learning (MacIntyre et al., 2023). Some available studies compare anticipatory response patterns during sequence learning in child and adults with varying musical experience, and do not isolate the effect of music training from age-related cognitive development. For example, Palmer and Pfordresher (2003) contrasted child novices with adult musicians, but the design did not control for age, making it difficult to attribute observed group differences in anticipatory response patterns specifically to training rather than developmental differences. On the other hand, Drake and Palmer (2000) show that increases in anticipatory planning are closely linked to both age and accumulated musical experience, with age often exerting a stronger influence than formal training alone. Our findings are in line with a recent study that showed more anticipatory eye movements in musicians when sequences were learned via eye movements (Schwizer Ashkenazi et al., 2022). Extending this line of work, we provide clear evidence for an anticipatory mode of responding during sequence learning that is augmented by practice and by musical experience.

Exactly what mechanisms drive this pattern of anticipatory responding in those with music training? One possibility is that music training results in a generalised improvement in predictive processing. Music training might improve the brain’s predictive processing of exogenous stimuli, such that musicians are better than non-musicians at predicting upcoming items in auditory sequences, even when those sequences have complex or high-order structures (Pesnot Lerousseau & Schön, 2021). Another possibility is that musicians are faster and better at integrating multisensory information (Landry & Champoux, 2017), which would allow them to form more robust audio-visual-motor representations during acquisition.

## Conclusion

In conclusion, our findings demonstrate that the stability of a new sensorimotor memory is contingent on the relevance of the sensory modality being challenged. The acquisition process itself is a strategic shift toward prediction, a shift that is accelerated in musicians. We speculate that this predictive pattern of behaviour results from an enhancement of latent processes with musical training, such as multisensory integration and/or predictive temporal attention.

## Declaration of generative AI and AI-assisted technologies in the manuscript preparation process

During the preparation of this work, the authors used ChatGPT-4o to improve readability and language of their original text. After using this tool, they reviewed and edited the content as needed and take full responsibility for the content of the publication.

